# Unified Sampling and Ranking for Protein Docking with DFMDock

**DOI:** 10.1101/2024.09.27.615401

**Authors:** Lee-Shin Chu, Sudeep Sarma, Da Xu, Jeffrey J. Gray

## Abstract

Recent diffusion-based approaches to protein-protein docking typically decouple structure generation from decoy ranking. We introduce **DFMDock** (Denoising Force Matching for Docking), a unified diffusion model that integrates generative sampling and energy-based ranking through physically motivated supervision. DFMDock predicts both denoising forces and a scalar energy, trained using force matching and energy contrastive objectives. The predicted forces guide the reverse diffusion process, while the energy enables decoy ranking without relying on a separately trained confidence model. On the Docking Benchmark 5, DFMDock achieves a 32.8% Oracle success rate and 5.3% Top-1 success rate, outperforming DiffDock-PP (16.2% and 4.3%, respectively). Unlike co-folding models, DFMDock does not require MSAs and generalizes to unseen targets. In decoy ranking, its learned energy function outperforms Rosetta energy and model-derived confidence scores, producing funnel-shaped energy landscapes enriched for near-native structures. These results suggest DFMDock as an efficient and physically grounded approach to diffusion-based protein docking.

## 1 Introduction

Predicting the three-dimensional structure of a protein complex from its unbound components remains a fundamental challenge in structural biology. Classical docking pipelines approach this problem in two stages: generating candidate binding poses through global or local sampling, and evaluating them with physics-based or empirical scoring functions [1, 2]. Despite progress in improving sampling efficiency and energy accuracy, these methods often require extensive computational resources, especially when combined with backbone flexibility or all-atom refinement. Template-based docking can accelerate prediction for homologous pairs [3], but fails when no suitable templates are available.

Leveraging large databases of protein sequences [4] and structures [5], co-folding models like AlphaFold2 [6] and RoseTTAFold [7] treat structure prediction as a sequence-to-structure task using multiple sequence alignments (MSAs). While effective for stable complexes with deep sequence coverage, these models remain computationally expensive due to MSA generation and often underperform for transient interactions such as antibody-antigen binding [8]. AlphaRED [9] improves interface modeling by applying ReplicaDock-based flexible backbone refinement to AlphaFold-Multimer [10] predictions, yielding better accuracy for induced-fit cases. AlphaFold3 and its reimplementations [11–15] expand modeling capabilities to multi-molecular assemblies via a diffusion module, but still fail in a substantial fraction of antibody-antigen cases [16, 17], in part due to their reliance on MSAs.

To circumvent these limitations, regression-based deep-learning docking methods were developed to predict docking poses directly from unbound 3D structures. EquiDock [18] pioneered the use of equivariant neural networks for rigid-body docking. Subsequent models such as ElliDock [19], GeoDock [20], and DockGPT [21] build on this idea, with flexible backbones or AlphaFold-inspired designs. However, these models typically generate a single deterministic prediction, lack sampling diversity, and underperform compared to MSA-based methods.

Generative diffusion models offer a compelling alternative by modeling docking as a denoising trajectory from noise to the data distribution. DiffDock [22] introduced a diffusion-based framework for small molecule ligand docking, using score matching [23] to learn transformations in translation, rotation, and ligand torsion space. DiffDock-PP [24] extended this approach to rigid protein-protein docking. Additional variants such as DiffMaSIF [25] incorporate protein interface features, while LatentDock [26] applies diffusion in a latent space via a variational autoencoder [27], akin to Stable Diffusion [28]. These methods typically rely on a separately trained confidence model to rank decoys.

Energy-based models (EBMs) provide an alternative by learning an energy function whose gradient guides structure prediction. DockGame [29] combines supervised and self-supervised learning to train energy functions for docking, while EBMDock [30] employs statistical potentials and Langevin dynamics to guide sampling. Arts et al. [31] used diffusion-based force fields for molecular dynamics by matching energy gradients to denoising forces. DSMBind [32] applies this idea to protein docking, showing that the learned energy aligns well with binding affinity.

Here we introduce DFMDock, a diffusion-based framework for rigid protein–protein docking that integrates structure generation and pose ranking through a learned energy function. The model is trained to predict translational and rotational forces using physically motivated objectives, including force matching and contrastive learning, without relying on multiple sequence alignments. This work addresses several open questions: Can a diffusion-based generative model learn an energy landscape that effectively distinguishes near-native from incorrect poses? Can it rank its own samples without a separate confidence model? And can it generalize to unseen protein pairs while producing diverse and physically plausible docking decoys? These questions motivate the design and evaluation of DFMDock.

**Figure 1.**
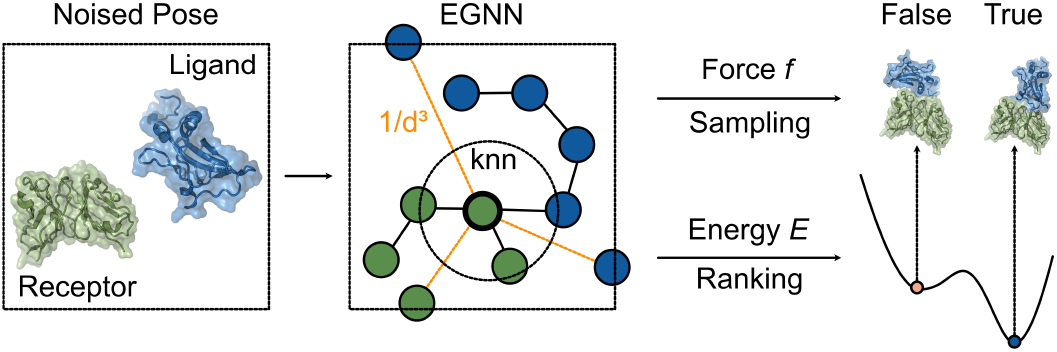
DFMDock model overview.

## 2 Methods

### 2.1 Learning energy and force in a diffusion model

According to statistical physics, a docking pose *x* is a random state sampled from the Boltzmann distribution 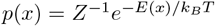, where *E*(*x*) is the energy, *k*_*B*_ is the Boltzmann constant, *T* is the temperature, and 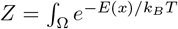 is the partition function. For rigid docking between a “receptor” protein and a “ligand” protein, the search space Ω spans all possible rigid-body translations and rotations of the ligand, with the receptor fixed. The per-residue forces on the ligand are given by:

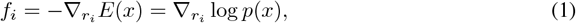

where *r*_*i*_ are the *C*_*α*_ coordinates of the ligand residues relative to their center. The model includes *N* and *C* atoms to compute orientation features in the input, and it captures translational forces on each residue (acting at the *C*_*α*_ atom) but not intra-residue torques. The model objective is to learn *f*_*i*_ and *E*(*x*), using *f*_*i*_ for sampling docking poses and *E*(*x*) for ranking.

### 2.2 Model Architecture

To jointly predict the energy *E* of a protein-protein complex and the per-residue forces *f*_*i*_ on the ligand, we use an equivariant graph neural network (EGNN) [33]:

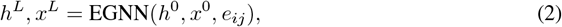

where *h* denotes the node embeddings, *x* the *C*_*α*_ coordinates, and *L* the number of graph network layers. The input node features *h*^0^ are constructed by concatenating a one-hot amino acid encoding, ESM-2 (650M) embeddings [34], and a binary indicator for receptor (0) or ligand (1). The edge features *e*_*ij*_ combine trRosetta-derived geometric priors [35] and relative positional encodings [10]. To model both local and long-range interactions, we construct hybrid graphs using 20 nearest neighbors and 40 additional edges sampled using inverse-cubic distance weighting [36].

The predicted energy is computed as a weighted sum over receptor-ligand residue pairs using a multi-layer perceptron *ϕ*_*E*_:

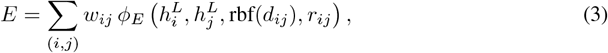

where *d*_*ij*_ is the Euclidean distance between residues *i* and *j*, and *r*_*ij*_ is the unit vector pointing from *i* to *j*. The term rbf(*d*_*ij*_) denotes a radial basis function encoding of the distance using 16 Gaussian kernels evenly spaced between 2 and 22 Å. The pairwise weights are defined as:

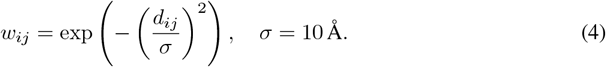

The predicted force on each ligand residue is defined as the predicted coordinate displacement after message passing:

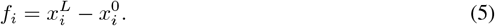

The total translational force on the ligand is:

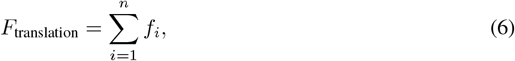

and the rotational force is:

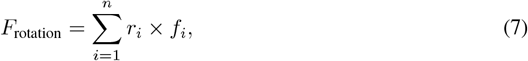

where *r*_*i*_ is the position of residue *i* relative to the ligand center of mass. As shown in the Appendix A, this formulation corresponds to computing gradients of the energy with respect to translational and rotational motion.

To improve numerical stability of force predictions [37], we normalize the translational and rotational forces and apply learned scaling using two MLPs *ϕ*translation and *ϕ*_rotation_, which are conditioned on the respective force magnitudes and the diffusion timestep *t*:

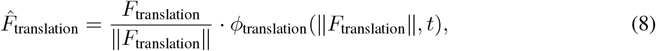

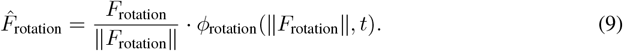

### 2.3 Loss Functions

We train the model using denoising force matching objectives for both translation and rotation. Here, the hatted variables 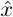 and 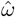 denote noised states sampled from the diffusion process, conditioned on the clean data states *x* and *ω*, respectively. The translation loss minimizes the difference between the predicted translational force and the score function (i.e., gradient of the log conditional density) of the noised translation:

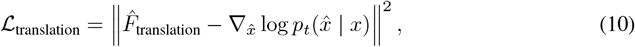

The rotation loss applies the same principle in the space of rotations:

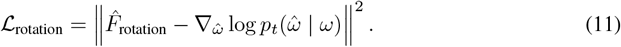

We use a Variance Exploding (VE) SDE [38] for both translation and rotation. Specifically, for translation, 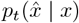 denotes a Gaussian distribution 𝒩 (*x, σ*_trans_(*t*)^2^**I**) in ℝ^3^; for rotation, 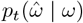 denotes an isotropic Gaussian 𝒩 (*ω, σ*_rot_(*t*)^2^**I**) in the Lie algebra of SO(3), following prior works [39– 41]. Both noise schedules are defined in Appendix B.

To align the predicted energy landscape with the predicted forces, we apply a force-matching loss [42], defined as:

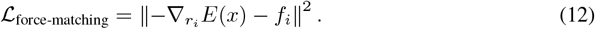

To ensure that native structures correspond to lower energies, we use a contrastive energy loss [43, 44]:

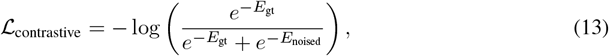

where *E*_gt_ and *E*_noised_ are the predicted energies for the ground-truth and noised structures, respectively.

We additionally apply a binary cross-entropy loss to classify interface residues:

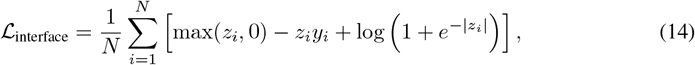

where *z*_*i*_ is the logits for residue *i*, obtained by applying a learned linear projection to its final-layer embedding 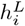, and *y*_*i*_ ∈ {0, 1} is the ground-truth interface label defined by any inter-chain *C*_*α*_ − *C*_*α*_ distance within 10 Å.

The total loss is a sum of all components:

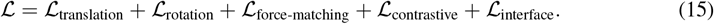

### 2.4 Confidence Model

To compare energy-based and confidence-base rankings, we trained a separate confidence model with the same architecture as the main model. The confidence model inputs poses generated by running 20 reverse diffusion steps from the trained diffusion model and outputs a scalar logit indicating the likelihood of the pose being near-native. Following the objective of DiffDock-PP [24], the confidence model is trained using binary cross-entropy loss, with poses labeled positive if ligand RMSD < 5.0 Å and negative otherwise.

### 2.5 Inference

During inference, DFMDock generates docking poses using a denoising diffusion process conditioned on randomized initial placements of the ligand. For each protein-protein target, the ligand is initialized with a random translation sampled from a Gaussian distribution with zero mean and standard deviation 30 Å, and a random orientation uniformly sampled from SO(3). A sequence of 40 reverse diffusion steps is then applied.

At each step, the ligand is updated using the predicted translational and rotational forces. The translational update shifts the ligand coordinates, while the rotational update applies an angular displacement using the exponential map on SO(3) around ligand’s current center of mass. This process iteratively refines the ligand pose toward a plausible binding configuration.

## 3 Experiments

### 3.1 Data

DFMDock was trained on DIPS-hetero, a subset of the DIPS dataset [45, 46], consisting of approximately 11,000 heterodimeric protein complexes. Evaluation was conducted on the Docking Benchmark 5 (DB5) dataset [47], which includes 253 protein-protein targets spanning a variety of complex types.

### 3.2 Comparisons

We compared DFMDock against two recent docking methods:

- **DiffDock-PP** [24], a diffusion-based model for protein-protein docking. We followed the instructions at https://github.com/ketatam/DiffDock-PP.
- **Boltz-1** [13], an open-source reimplementation of AlphaFold3 that predicts multimeric structures from sequences and ranks candidates using a learned confidence model. We ran Boltz-1 in both standard and No-MSA modes, following the instructions at https://github.com/jwohlwend/boltz/tree/main.

All methods were evaluated on the same DB5 targets using their publicly released inference configurations.

### 3.3 Evaluation Protocol

Docking accuracy was measured using the DockQ score [48], with thresholds of 0.23, 0.49, and 0.80 corresponding to acceptable, medium, and high-quality predictions, respectively, following CAPRI community custom [49]. We report Oracle and Top-*k* success rates, defined as the percentage of targets for which at least one decoy of acceptable quality is found anywhere in the candidate set (Oracle) or within the top *k* predictions ranked with the DFMDock energy or the confidence metrics of DiffDock-PP or Boltz-1 (Top-*k*).

### 3.4 Training Setup

DFMDock was trained for 50 epochs using the Adam optimizer with a learning rate of 1 *×* 10^−4^ and an effective batch size of 4. Training was performed using PyTorch’s Distributed Data Parallel (DDP) across four NVIDIA A100 GPUs, and completed in approximately 8 hours.

## 4 Results

### 4.1 DFMDock outperforms DiffDock-PP and Boltz-1 (No MSA)

We compared DFMDock to DiffDock-PP, Boltz-1, and Boltz-1 (No MSA) on the Docking Benchmark 5 (DB5). DFMDock and DiffDock-PP each generated 40 samples per target using 40 diffusion steps from randomized initial poses, while Boltz-1 and Boltz-1 (No MSA) generated 5 samples per target, ranked by their confidence scores. DFMDock ranked samples using its learned energy function, and DiffDock-PP used its internal confidence model. The results for DFMDock and DiffDock-PP were averaged over three independent runs with different random seeds.

As shown in Figure 2, DFMDock outperforms DiffDock-PP across all evaluation metrics. In the Oracle setting, which considers whether any sampled pose is acceptable, DFMDock achieved a success rate of 32.8%, compared to 16.2% for DiffDock-PP. For Top-5 and Top-1 rankings, which assess the ability to prioritize near-native poses, DFMDock achieved 13.8% and 5.3% success, compared to 7.9% and 4.3% for DiffDock-PP. These results indicate that DFMDock is more effective both at generating high-quality poses and at identifying them through its energy-based ranking.

**Figure 2.**
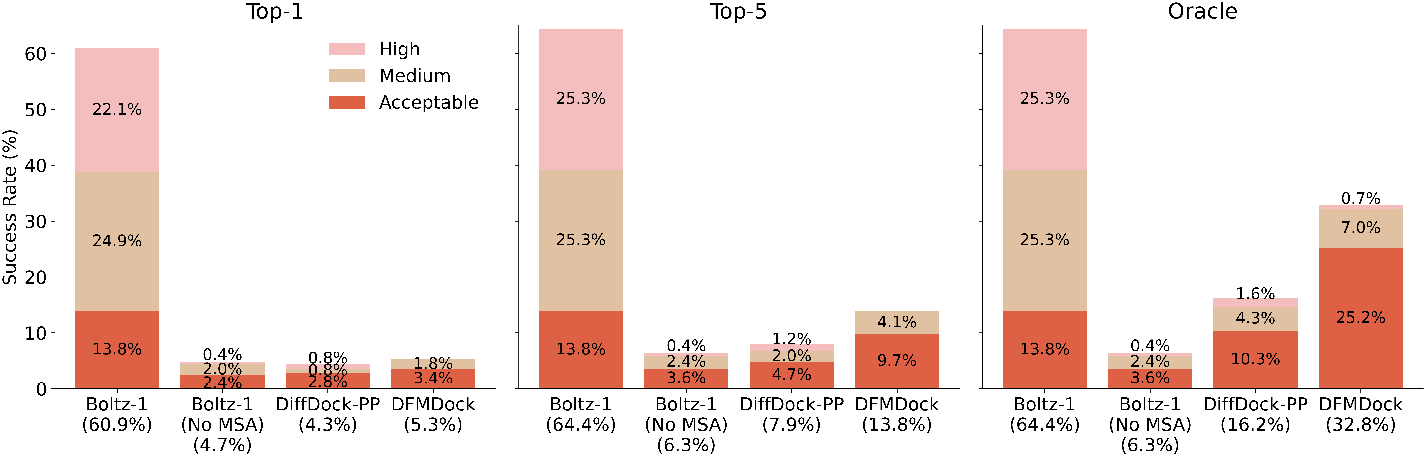
Success rates of Boltz-1, Boltz-1 (No MSA), DiffDock-PP, and DFMDock on the Docking Benchmark 5 (*N* = 253) under Top-1, Top-5, and Oracle evaluation settings. Bars are segmented by DockQ [48] thresholds: Acceptable (0.23 ≤ DockQ *<* 0.49), Medium (0.49 ≤ DockQ *<* 0.80), and High (DockQ ≥ 0.80).

Boltz-1 achieved the highest overall success rates, but its performance benefits from multiple sequence alignments (MSAs) and partial overlap between DB5 and its training data. In contrast, DFMDock requires no MSAs and was trained on a non-overlapping dataset. Indeed, removing the MSA from Boltz-1 results in much lower success rates. The lower performance of DiffDock-PP on DB5, relative to its original evaluation on DIPS [24], may reflect test-set leakage [20, 50], suggesting stronger generalization by DFMDock.

### 4.2 DFMDock energy exhibits funnel-like behavior and outperforms Rosetta energy and confidence scores in decoy ranking

To assess the ranking capability of DFMDock’s learned energy function, we compared it against two baselines: the Rosetta energy function [51] and a trained confidence model. Figure 3 shows results for the USP14–ubiquitin aldehyde complex (PDB ID: 2AYO). DFMDock energy displays a funnel-shaped landscape, where lower energy values cluster near low interface RMSD and correspond to high DockQ decoys. That is, the model assigns more favorable scores to near-native structures. In comparison, Rosetta energy shows a weaker funnel, and the confidence model exhibits little separation across RMSD values. These patterns are reflected in the *P*_near_ metric [52], which quantifies the quality of funnel-like energy landscapes. Energy funnels and *P*_near_ values for the remaining 24 targets and the summary statistics are shown in Supplementary Figure S1 and Table S1.

**Figure 3.**
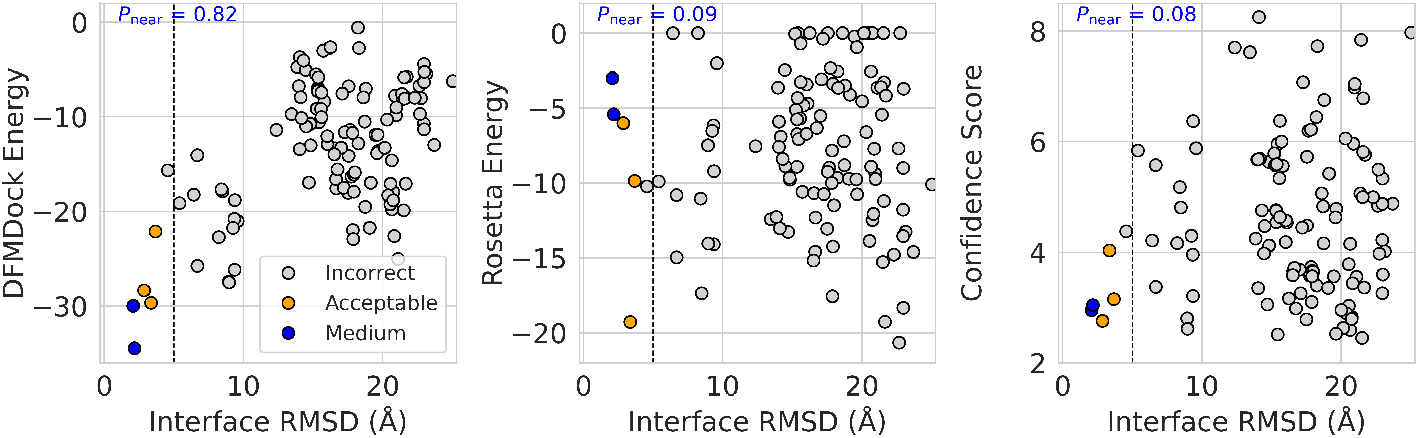
Comparison of decoy ranking metrics using DFMDock energy, Rosetta energy, and model-derived confidence scores. A case study on the USP14–ubiquitin aldehyde complex (PDB ID: 2AYO) showing interface RMSD versus ranking scores. Each point represents a decoy, colored by DockQ-based structural quality: gray (incorrect, DockQ *<* 0.23), orange (acceptable, 0.23 ≤ DockQ *<* 0.49), and blue (medium, 0.49 ≤ DockQ *<* 0.80). The vertical dashed line marks the 5 Å threshold for acceptable predictions. The *P*_near_ value shown in each panel quantifies the degree of energy funneling (higher is better).

Figure 4a shows top-*k* success rates across 25 DB5 targets. DFMDock outperforms both baselines in identifying acceptable-quality decoys among the top-*k* predictions. Success rate is defined as the proportion of targets with at least one acceptable structure in the top *k* decoys. Results are averaged over three runs with different random seeds; error bars show standard deviation. The largest improvements occur in the top-1 and top-2 settings. Differences diminish beyond top-3, but DFMDock maintains higher overall performance.

**Figure 4.**
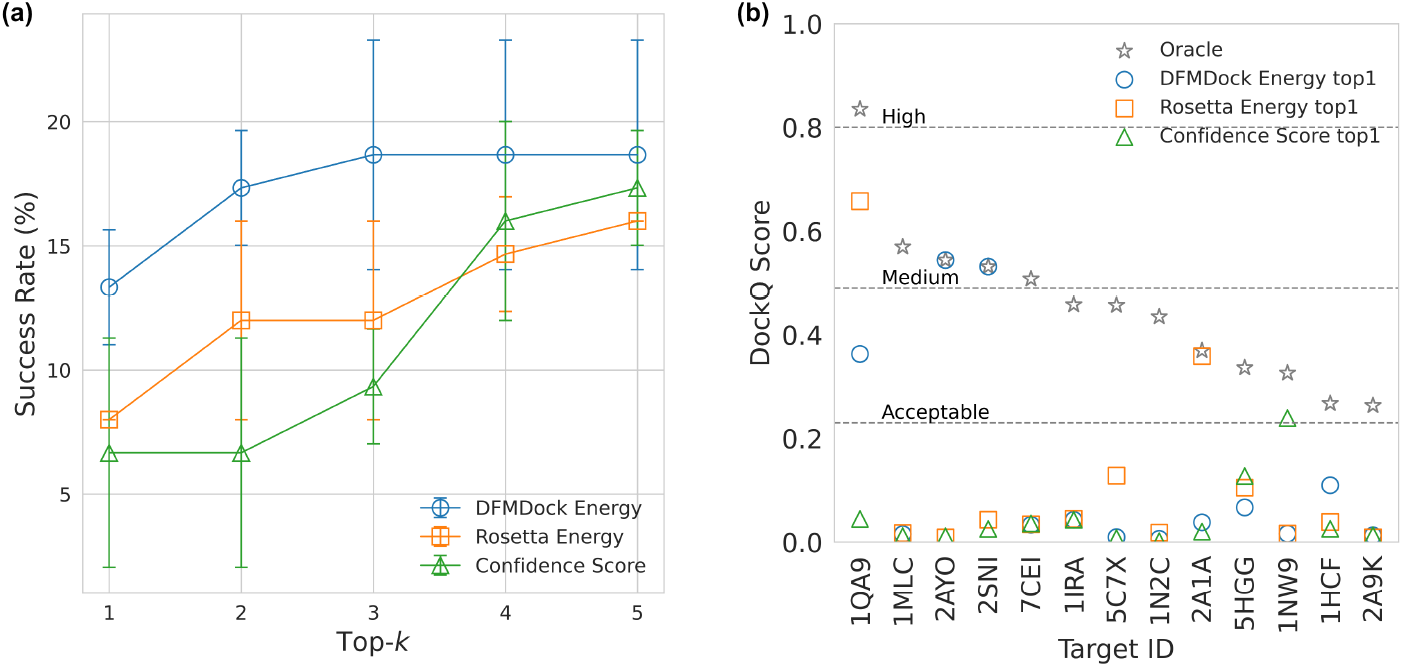
Comparison of decoy ranking metrics using DFMDock energy, Rosetta energy, and model-derived confidence scores. **(a)** Top-*k* success rates on the DB5 subset (*N* = 25), defined as the percentage of targets with at least one acceptable-quality decoy (DockQ ≥ 0.23) among the top *k* predictions. Error bars indicate standard deviation across three independent runs. **(b)** Per-target comparison of the DockQ score of the best available decoy and the top-ranked decoy under each metric for the 13 targets with Oracle DockQ ≥ 0.23. Horizontal lines at DockQ = 0.23, 0.49, and 0.80 indicate thresholds for acceptable, medium, and high-quality predictions.

Figure 4b compares the DockQ score of the top-ranked decoy from each method to the best available decoy for each target. DFMDock energy more reliably selects near-native structures than either Rosetta or the confidence model. This is achieved without structure refinement, which is required for Rosetta. These results indicate that the DFMDock energy function supports more accurate decoy ranking without relying on refinement or a separate confidence model.

To complement the ranking results, Figure 5 shows a structural comparison of the top-ranked decoys for a subtilisin-chymotrypsin complex (PDB ID: 2SNI). The DFMDock Energy ranks a medium-quality decoy (DockQ = 0.53) as its top-1 prediction, while Rosetta energy and the confidence score select incorrect decoys with DockQ scores of 0.04 and 0.02, respectively. This example illustrates a case where The DFMDock energy more accurately identifies a near-native binding mode, consistent with its improved ranking performance across the DB5 test set.

**Figure 5.**
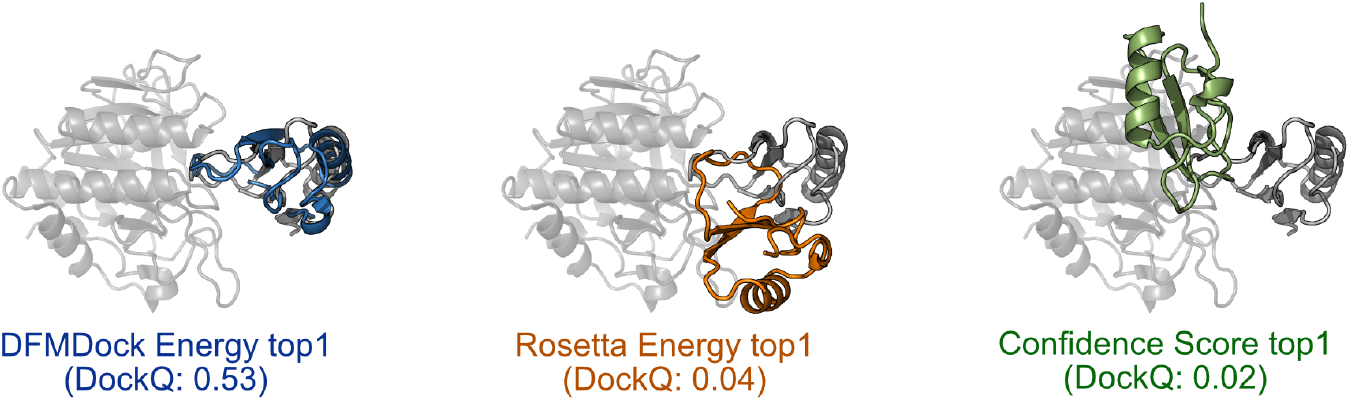
Top-ranked decoys for a subtilisin-chymotrypsin complex (PDB ID: 2SNI). Native receptor and ligand are shown in gray; predicted ligands are colored by ranking method. Left: Lowest DFMDock energy structure in blue; middle: Lowest Rosetta energy structure in orange; right: Best Confidence score structure in green.

### 4.3 Ablation study highlights the importance of force matching and contrastive loss

To assess the contribution of key training components, we conducted ablation experiments by removing force matching loss (No FM), contrastive loss (No CL), or both (No FMCL) from the DFMDock model. As shown in Figure 6, which summarizes results on the same 25-target subset, removing either component resulted in reduced performance across all evaluation criteria, including Top-1, Top-5, and Oracle success rates. Reported values are averaged over three independent runs with different random seeds.

**Figure 6.**
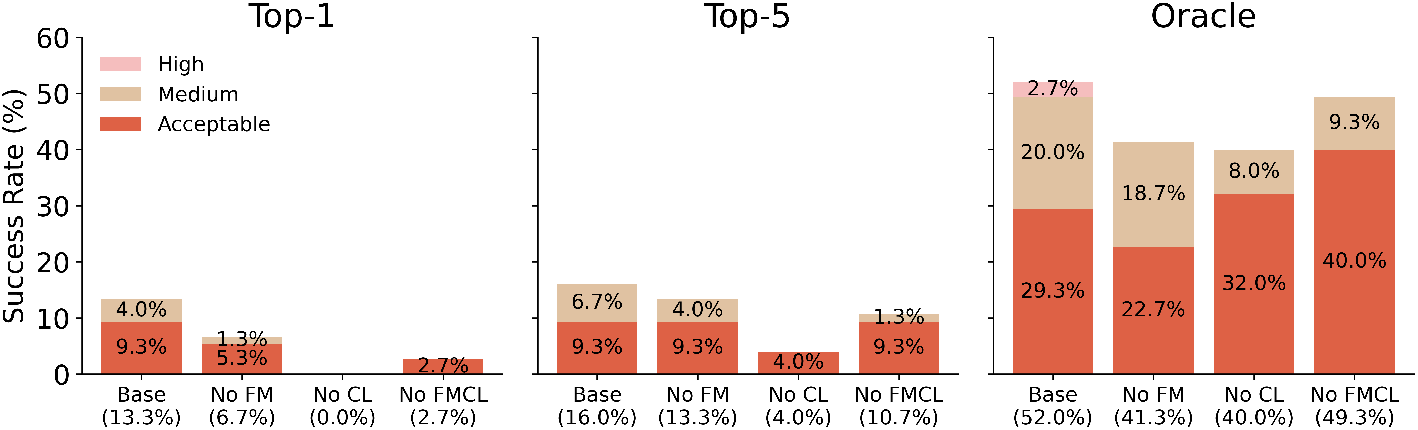
Ablation study of model components evaluated using Top-1, Top-5, and Oracle success rates on the DB5 subset (*N* = 25). Each stacked bar represents the performance of a model variant with one component removed: Force Matching (No FM), Contrastive loss (No CL), or both No FMCL. Bars are segmented by DockQ thresholds into Acceptable (0.23 ≤ DockQ *<* 0.49), Medium (0.49 ≤ DockQ *<* 0.80), and High (DockQ ≥ 0.80).

Excluding contrastive loss led to the most pronounced decline. In this setting, the model failed to produce any acceptable Top-1 predictions (0%), and Top-5 success dropped to 4.0%, compared to 13.3% and 16.0% in the full model. These results indicate that contrastive supervision is critical for guiding the model to generate and correctly rank near-native structures. Removing force matching also reduced performance, with Top-1 success falling to 6.7% and Oracle success to 41.3%, showing that force-based supervision improves both the quality of sampled decoys and their ranking. The model trained without both components (No FMCL) outperformed the No CL variant across all metrics but remained below the base and No FM models. In the Oracle setting, No FMCL achieved higher overall success than No FM (49.3% vs. 41.3%) but yielded fewer medium-quality predictions (9.3% vs. 18.7%), suggesting that while more acceptable decoys were found, they were of lower quality. These results show that both training signals are necessary to support accurate ranking and robust sampling.

## 5 Discussion and Conclusion

These results suggest that diffusion-based docking can be improved by incorporating an energy objective in the model, particularly for approaches that do not use multiple sequence alignments (MSAs) or computationally intensive refinement. DFMDock learns an energy landscape using force matching and contrastive supervision, without requiring task-specific heuristics or pretrained confidence models. Compared to DiffDock-PP, the model generalizes better to unseen targets, supporting the effectiveness of energy-based supervision in the absence of MSAs.

DFMDock integrates force matching and contrastive loss to align its generative process with physical and structural constraints. Force supervision encourages coordinate updates consistent with molecular interactions, while the contrastive objective ensures that near-native decoys are assigned lower energy than perturbed or incorrect structures. Ablation experiments indicate that both components are necessary for optimal performance. The resulting energy function enables effective ranking directly from generated structures without refinement or confidence models, simplifying the inference pipeline.

Compared to existing models, DFMDock offers a flexible and generative approach. DSMBind [32] focuses on binding affinity prediction and does not generate docked structures, making it unsuitable for docking. EBMDock [30] and DockGame [29] report per-target metrics such as DockQ, RMSD, and TM-score averaged over benchmark sets but do not provide target-level success rates, limiting direct comparison to DFMDock’s top-*k* success metrics. For example, the EBMDock paper reports an average DockQ of 0.05 on DB5, similar to DiffDock-PP’s 0.04, indicating comparably low accuracy across the benchmark. GeoDock [20] and DockGPT [21] take a regression-based approach that directly predicts complex structures without sampling, which limits their ability to represent multiple plausible docking poses.

DFMDock is less accurate than co-folding-based methods, which leverage MSA-derived features and full-sequence context to capture co-evolutionary constraints. These models can implicitly account for some structural flexibility but are not explicitly designed to model induced fit or large conformational changes. Operating without MSAs and assuming rigid backbones, DFMDock is limited in resolving complex interfaces. However, its computational efficiency and independence from external sequence databases make it a practical alternative to MSA-dependent methods.

The force matching framework could extend naturally to all-atom diffusion models (e.g., AF3 or Boltz), where energy-based supervision may improve fine-grained accuracy. Incorporating flexible backbone and side-chain sampling could further enhance performance. DFMDock provides a foundation for integrating physical priors into generative models for macromolecular docking.

## Acknowledgements

This work was supported by National Institutes of Health grant R35-GM141881 and by Moderna. Computational resources were provided by the Advanced Research Computing at Hopkins (ARCH). The authors thank Jeremias Sulam for valuable discussions and insightful feedback.

## Code Availability

The inference code, model weights, and test set are available at https://github.com/Graylab/DFMDock.

## A Gradient of energy with respect to a rotation vector

Consider a rigid body with *N* points labeled **r**_1_, **r**_2_, …, **r**_*N*_, where each **r**_*i*_ = **x**_*i*_ − **c** represents the position of point *i* relative to the center of mass **c**. When the rigid body undergoes a small rotation *dω* through its center of mass, where *ω* is a rotation vector, the displacement *d***r**_*i*_ of each point **r**_*i*_ is given by:

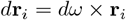

The energy *E* of the system depends on the positions **r**_*i*_. The change in energy *dE* due to small displacements *d***r**_*i*_ of the points is:

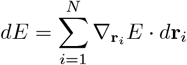

where 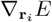 is the gradient of the energy with respect to the position of point **r**_*i*_.

Substituting the expression for the displacement *d***r**_*i*_, we get:

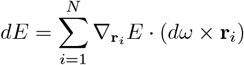

Using the scalar triple product identity:

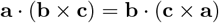

we can simplify the expression for *dE*:

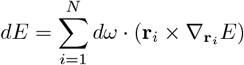

Thus, the gradient of energy with respect to the rotation vector *ω* is:

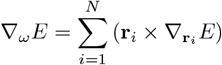

where **r**_*i*_ is the vector from the center of mass to the point **x**_*i*_, and 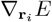 is the gradient of the energy with respect to the position of point **r**_*i*_.

## B Noise Schedules

### Translation noise schedule

The standard deviation used for translation noise is defined as

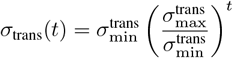

where 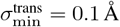 and 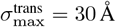

### Rotation noise schedule

The rotation noise schedule is given by

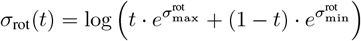

where 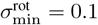 and 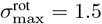

## Notes

### Competing Interest Statement

The authors have declared no competing interest.

### Summary of Updates

Sections updated Figures revised author affiliations updated

## References

[1] Ilya A Vakser. Protein-protein docking: From interaction to interactome. Biophysical journal, 107(8):1785–1793, 2014.

[2] Sheng-You Huang. Search strategies and evaluation in protein–protein docking: principles, advances and challenges. Drug Discovery Today, 19(8):1081–1096, 2014.

[3] Andras Szilagyi and Yang Zhang. Template-based structure modeling of protein–protein interactions. Current Opinion in Structural Biology, 24:10–23, 2014.

[4] The UniProt Consortium. Uniprot: The universal protein knowledgebase. Nucleic Acids Research, 46(5):2699–2699, 2018.

[5] Helen M Berman, Tammy Battistuz, Talapady N Bhat, Wolfgang F Bluhm, Philip E Bourne, Kyle Burkhardt, Zukang Feng, Gary L Gilliland, Lisa Iype, Shri Jain, et al. The protein data bank. Acta Crystallographica Section D: Biological Crystallography, 58(6):899–907, 2002.

[6] John Jumper, Richard Evans, Alexander Pritzel, Tim Green, Michael Figurnov, Olaf Ronneberger, Kathryn Tunyasuvunakool, Russ Bates, Augustin Žídek, Anna Potapenko, et al. Highly accurate protein structure prediction with AlphaFold. Nature, 596(7873):583–589, 2021.

[7] Minkyung Baek, Frank DiMaio, Ivan Anishchenko, Justas Dauparas, Sergey Ovchinnikov, Gyu Rie Lee, Jue Wang, Qian Cong, Lisa N Kinch, R Dustin Schaeffer, et al. Accurate prediction of protein structures and interactions using a three-track neural network. Science, 373(6557):871–876, 2021.

[8] Rui Yin, Brandon Y Feng, Amitabh Varshney, and Brian G Pierce. Benchmarking AlphaFold for protein complex modeling reveals accuracy determinants. Protein Science, 31(8):e4379, 2022.

[9] Ameya Harmalkar, Sergey Lyskov, and Jeffrey J Gray. Reliable protein-protein docking with AlphaFold, Rosetta, and replica-exchange. eLife, page RP94029, 2025.

[10] Richard Evans, Michael O’Neill, Alexander Pritzel, Natasha Antropova, Andrew Senior, Tim Green, Augustin Žídek, Russ Bates, Sam Blackwell, Jason Yim, et al. Protein complex prediction with AlphaFold-Multimer. biorxiv, pages 2021–10, 2021.

[11] Josh Abramson, Jonas Adler, Jack Dunger, Richard Evans, Tim Green, Alexander Pritzel, Olaf Ronneberger, Lindsay Willmore, Andrew J Ballard, Joshua Bambrick, et al. Accurate structure prediction of biomolecular interactions with alphafold 3. Nature, pages 1–3, 2024.

[12] Chai Discovery team, Jacques Boitreaud, Jack Dent, Matthew McPartlon, Joshua Meier, Vinicius Reis, Alex Rogozhonikov, and Kevin Wu. Chai-1: Decoding the molecular interactions of life. BioRxiv, pages 2024–10, 2024.

[13] Jeremy Wohlwend, Gabriele Corso, Saro Passaro, Mateo Reveiz, Ken Leidal, Wojtek Swiderski, Tally Portnoi, Itamar Chinn, Jacob Silterra, Tommi Jaakkola, et al. Boltz-1: Democratizing biomolecular interaction modeling. bioRxiv, pages 2024–11, 2024.

[14] Lihang Liu, Shanzhuo Zhang, Yang Xue, Xianbin Ye, Kunrui Zhu, Yuxin Li, Yang Liu, Jie Gao, Wenlai Zhao, Hongkun Yu, et al. Technical report of helixfold3 for biomolecular structure prediction. arXiv preprint 2408.16975, 2024.

[15] ByteDance AML AI4Science Team, Xinshi Chen, Yuxuan Zhang, Chan Lu, Wenzhi Ma, Jiaqi Guan, Chengyue Gong, Jincai Yang, Hanyu Zhang, Ke Zhang, et al. Protenix-advancing structure prediction through a comprehensive alphafold3 reproduction. bioRxiv, pages 2025–01, 2025.

[16] Fatima N Hitawala and Jeffrey J Gray. What does alphafold3 learn about antigen and nanobody docking, and what remains unsolved? bioRxiv, pages 2024–09, 2024.

[17] Sheng Xu, Qiantai Feng, Lifeng Qiao, Hao Wu, Tao Shen, Yu Cheng, Shuangjia Zheng, and Siqi Sun. Foldbench: An all-atom benchmark for biomolecular structure prediction. bioRxiv, pages 2025–05, 2025.

[18] Octavian-Eugen Ganea, Xinyuan Huang, Charlotte Bunne, Yatao Bian, Regina Barzilay, Tommi Jaakkola, and Andreas Krause. Independent se (3)-equivariant models for end-to-end rigid protein docking. arXiv preprint 2111.07786, 2021.

[19] Ziyang Yu, Wenbing Huang, and Yang Liu. Rigid protein-protein docking via equivariant elliptic-paraboloid interface prediction. arXiv preprint 2401.08986, 2024.

[20] Lee-Shin Chu, Jeffrey A Ruffolo, Ameya Harmalkar, and Jeffrey J Gray. Flexible protein– protein docking with a multitrack iterative transformer. Protein Science, 33(2):e4862, 2024.

[21] Matt McPartlon and Jinbo Xu. Deep learning for flexible and site-specific protein docking and design. BioRxiv, pages 2023–04, 2023.

[22] Gabriele Corso, Hannes Stärk, Bowen Jing, Regina Barzilay, and Tommi Jaakkola. Diffdock: Diffusion steps, twists, and turns for molecular docking. arXiv preprint 2210.01776, 2022.

[23] Pascal Vincent. A connection between score matching and denoising autoencoders. Neural computation, 23(7):1661–1674, 2011.

[24] Mohamed Amine Ketata, Cedrik Laue, Ruslan Mammadov, Hannes Stärk, Menghua Wu, Gabriele Corso, Céline Marquet, Regina Barzilay, and Tommi S Jaakkola. Diffdock-pp: Rigid protein-protein docking with diffusion models. arXiv preprint 2304.03889, 2023.

[25] Freyr Sverrisson, Mehmet Akdel, Dylan Abramson, Jean Feydy, Alexander Goncearenco, Yusuf Adeshina, Daniel Kovtun, Céline Marquet, Xuejin Zhang, David Baugher, et al. Diffmasif: Surface-based protein-protein docking with diffusion models. In Machine Learning in Structural Biology workshop at NeurIPS 2023, 2023.

[26] Matt McPartlon, Céline Marquet, Tomas Geffner, Daniel Kovtun, Alexander Goncearenco, Zachary Carpenter, Luca Naef, Michael Bronstein, and Jinbo Xu. Latentdock: Protein-protein docking with latent diffusion. MLSB, 2023.

[27] Diederik P Kingma. Auto-encoding variational bayes. arXiv preprint 1312.6114, 2013.

[28] Robin Rombach, Andreas Blattmann, Dominik Lorenz, Patrick Esser, and Björn Ommer. High-resolution image synthesis with latent diffusion models. In Proceedings of the IEEE/CVF conference on computer vision and pattern recognition, pages 10684–10695, 2022.

[29] Vignesh Ram Somnath, Pier Giuseppe Sessa, Maria Rodriguez Martinez, and Andreas Krause. Dockgame: Cooperative games for multimeric rigid protein docking. arXiv preprint 2310.06177, 2023.

[30] Huaijin Wu, Wei Liu, Yatao Bian, Jiaxiang Wu, Nianzu Yang, and Junchi Yan. Ebmdock: Neural probabilistic protein-protein docking via a differentiable energy model. In The Twelfth International Conference on Learning Representations, 2024.

[31] Marloes Arts, Victor Garcia Satorras, Chin-Wei Huang, Daniel Zugner, Marco Federici, Cecilia Clementi, Frank Noé, Robert Pinsler, and Rianne van den Berg. Two for one: Diffusion models and force fields for coarse-grained molecular dynamics. Journal of Chemical Theory and Computation, 19(18):6151–6159, 2023.

[32] Wengong Jin, Xun Chen, Amrita Vetticaden, Siranush Sarzikova, Raktima Raychowdhury, Caroline Uhler, and Nir Hacohen. Dsmbind: Se (3) denoising score matching for unsupervised binding energy prediction and nanobody design. bioRxiv, pages 2023–12, 2023.

[33] Victor Garcia Satorras, Emiel Hoogeboom, and Max Welling. E (n) equivariant graph neural networks. In International conference on machine learning, pages 9323–9332. PMLR, 2021.

[34] Zeming Lin, Halil Akin, Roshan Rao, Brian Hie, Zhongkai Zhu, Wenting Lu, Nikita Smetanin, Robert Verkuil, Ori Kabeli, Yaniv Shmueli, et al. Evolutionary-scale prediction of atomic-level protein structure with a language model. Science, 379(6637):1123–1130, 2023.

[35] Jianyi Yang, Ivan Anishchenko, Hahnbeom Park, Zhenling Peng, Sergey Ovchinnikov, and David Baker. Improved protein structure prediction using predicted interresidue orientations. Proceedings of the National Academy of Sciences, 117(3):1496–1503, 2020.

[36] John B Ingraham, Max Baranov, Zak Costello, Karl W Barber, Wujie Wang, Ahmed Ismail, Vincent Frappier, Dana M Lord, Christopher Ng-Thow-Hing, Erik R Van Vlack, et al. Illuminating protein space with a programmable generative model. Nature, 623(7989):1070–1078, 2023.

[37] Matthew Masters, Amr Mahmoud, and Markus Lill. Fusiondock: Physics-informed diffusion model for molecular docking. In ICML2023 CompBio Workshop, 2023.

[38] Yang Song, Jascha Sohl-Dickstein, Diederik P Kingma, Abhishek Kumar, Stefano Ermon, and Ben Poole. Score-based generative modeling through stochastic differential equations. arXiv preprint 2011.13456, 2020.

[39] Adam Leach, Sebastian M Schmon, Matteo T Degiacomi, and Chris G Willcocks. Denoising diffusion probabilistic models on so (3) for rotational alignment. ICLR2022 GTRL Workshop, 2022.

[40] Jason Yim, Brian L Trippe, Valentin De Bortoli, Emile Mathieu, Arnaud Doucet, Regina Barzilay, and Tommi Jaakkola. Se (3) diffusion model with application to protein backbone generation. arXiv preprint 2302.02277, 2023.

[41] Yesukhei Jagvaral, Francois Lanusse, and Rachel Mandelbaum. Unified framework for diffusion generative models in so (3): applications in computer vision and astrophysics. In Proceedings of the AAAI Conference on Artificial Intelligence, 2024.

[42] Alexandre Agm Duval, Victor Schmidt, Alex Hernández-Garcia, Santiago Miret, Fragkiskos D Malliaros, Yoshua Bengio, and David Rolnick. Faenet: Frame averaging equivariant gnn for materials modeling. In International Conference on Machine Learning, pages 9013–9033. PMLR, 2023.

[43] Yilun Du, Jiayuan Mao, and Joshua B Tenenbaum. Learning iterative reasoning through energy diffusion. arXiv preprint 2406.11179, 2024.

[44] Changsoo Lee, Jonghun Won, Seongok Ryu, Jinsol Yang, Nuri Jung, Hahnbeom Park, and Chaok Seok. Galaxydock-dl: Protein–ligand docking by global optimization and neural network energy. Journal of Chemical Theory and Computation, 2024.

[45] RJL Townshend, R Bedi, PA Suriana, and RO Dror. End-to-end learning on 3d protein structure for interface prediction. arxiv. 1807.01297, 2018.

[46] Alex Morehead, Chen Chen, Ada Sedova, and Jianlin Cheng. Dips-plus: The enhanced database of interacting protein structures for interface prediction. Scientific Data, 10(1):509, 2023.

[47] Thom Vreven, Iain H Moal, Anna Vangone, Brian G Pierce, Panagiotis L Kastritis, Mieczyslaw Torchala, Raphael Chaleil, Brian Jiménez-García, Paul A Bates, Juan Fernandez-Recio, et al. Updates to the integrated protein–protein interaction benchmarks: docking benchmark version 5 and affinity benchmark version 2. Journal of Molecular Biology, 427(19):3031–3041, 2015.

[48] Sankar Basu and Björn Wallner. Dockq: a quality measure for protein-protein docking models. PloS One, 11(8):e0161879, 2016.

[49] Joël Janin, Kim Henrick, John Moult, Lynn Ten Eyck, Michael JE Sternberg, Sandor Vajda, Ilya Vakser, and Shoshana J Wodak. Capri: a critical assessment of predicted interactions. Proteins: Structure, Function, and Bioinformatics, 52(1):2–9, 2003.

[50] Anton Bushuiev, Roman Bushuiev, Jiri Sedlar, Tomas Pluskal, Jiri Damborsky, Stanislav Mazurenko, and Josef Sivic. Revealing data leakage in protein interaction benchmarks. arXiv preprint 2404.10457, 2024.

[51] Rebecca F Alford, Andrew Leaver-Fay, Jeliazko R Jeliazkov, Matthew J O’Meara, Frank P DiMaio, Hahnbeom Park, Maxim V Shapovalov, P Douglas Renfrew, Vikram K Mulligan, Kalli Kappel, et al. The rosetta all-atom energy function for macromolecular modeling and design. Journal of Chemical Theory and Computation, 13(6):3031–3048, 2017.

[52] Gaurav Bhardwaj, Vikram Khipple Mulligan, Christopher D Bahl, Jason M Gilmore, Peta J Harvey, Olivier Cheneval, Garry W Buchko, Surya VSRK Pulavarti, Quentin Kaas, Alexander Eletsky, et al. Accurate de novo design of hyperstable constrained peptides. Nature, 538(7625): 329–335, 2016.

