## Supplemental Table 1 and Figure 1 for "Unified Sampling and Ranking for Protein Docking with DFMDock"

**Lee-Shin Chu**[1] **Sudeep Sarma**[1] **Da Xu**[1] **Jeffrey J. Gray**[1,2,3,4]
[1]Department of Chemical and Biomolecular Engineering
[2]Program in Molecular Biophysics
[3]Data Science and AI Institute
[4]The Sidney Kimmel Comprehensive Cancer Center
Johns Hopkins University, Baltimore, MD 21218, USA.

#### Supplementary Information

#### Calculation of $P_{\text{near}}$ score

For each target and each ranking metric, we calculated the $P_{\text{near}}$ score [1]:

$$P_{\text{near}} = \frac{\sum_{i=1}^{N} e^{-E_i/kT} e^{-(\text{iRMSD}_i/\lambda)^2}}{\sum_{j=1}^{N} e^{-E_j/kT}}.$$

The $P_{\text{near}}$ score ranges from 0 (poor funneling) to 1 (sharp funnel with near-native low-energy decoys). Here, $E_i$ is the predicted energy or score of decoy $i$, and iRMSD$_i$ is its interface RMSD to the native structure. We used $\lambda = 5$ Å to reflect the docking criterion that interface RMSD below 5 Å is generally considered an acceptable prediction, whereas the original formulation of $P_{\text{near}}$ in protein design used $\lambda = 1$ Å for sub-angstrom accuracy. The inverse temperature parameter $1/kT$ was set to 1.0.

Table S1: Mean and standard deviation of the $P_{\text{near}}$ score for each ranking metric on the DB5 subset ($N = 25$).

| | DFMDock energy | Rosetta energy | Confidence score |
|---|---|---|---|
| Mean | 0.118 | 0.095 | 0.059 |
| Std | 0.247 | 0.213 | 0.085 |

Preprint. Under review.

### (a) DFMDock Energy

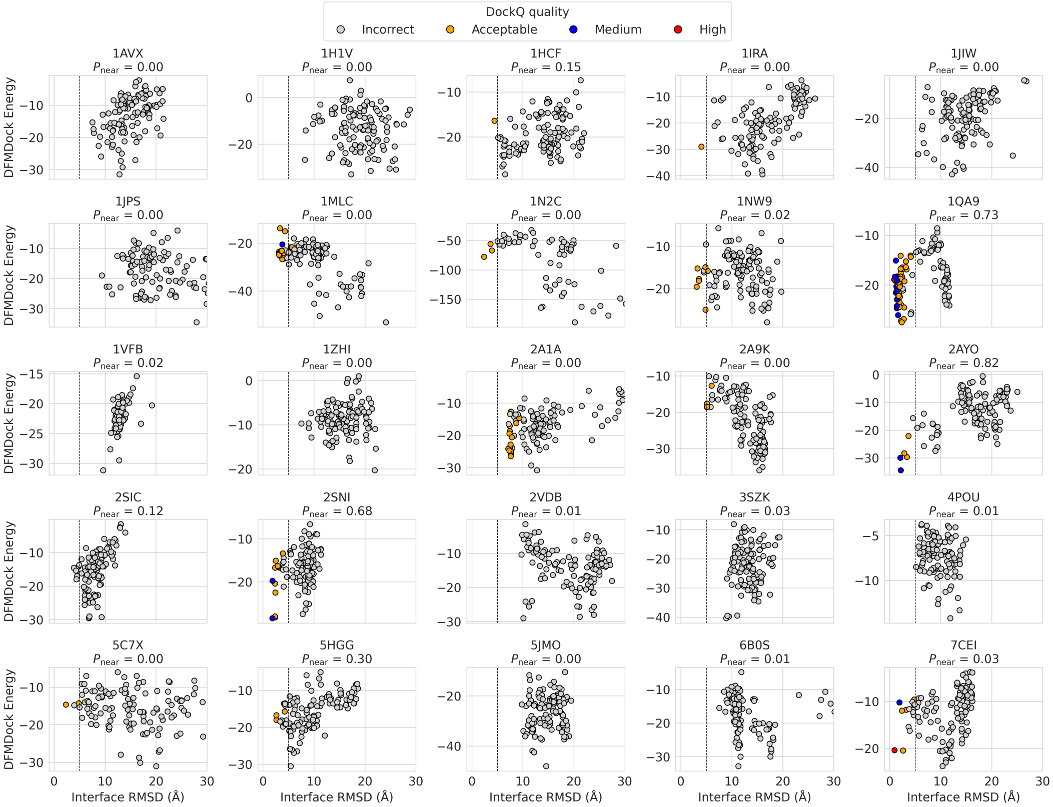


### (b) Rosetta Energy

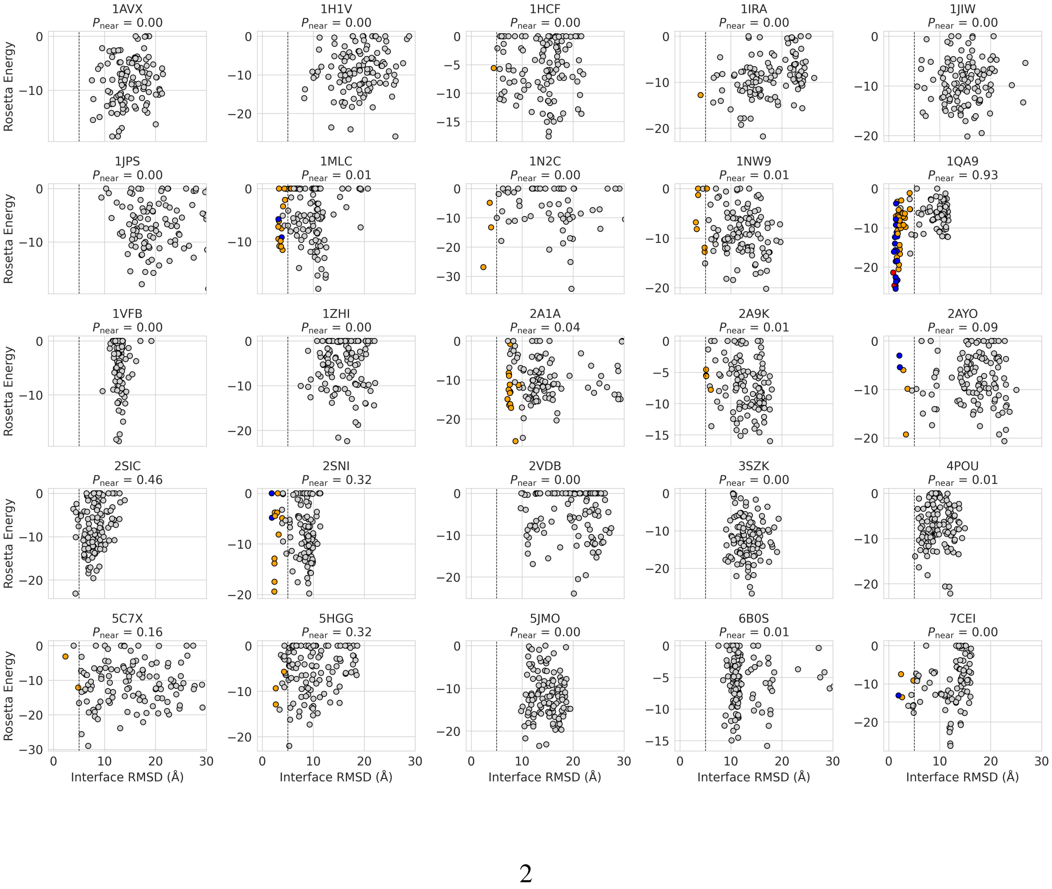

(c) Confidence Score

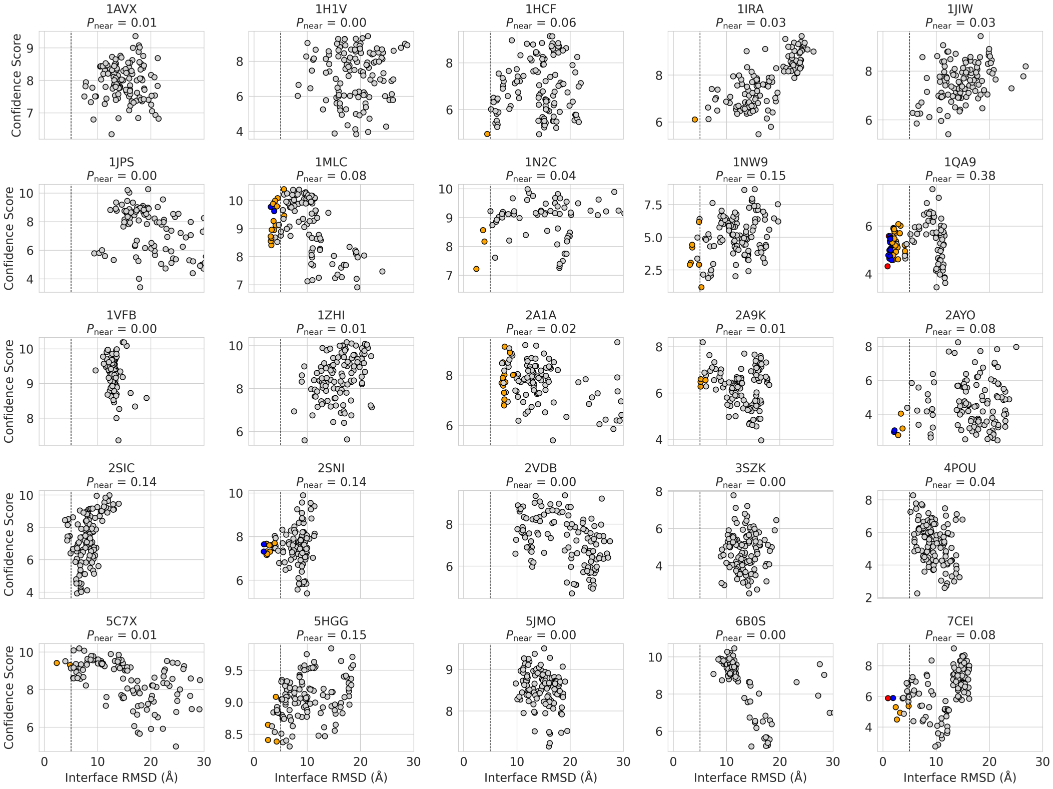


Figure S1: Interface RMSD versus ranking metrics for the DB5 subset ($N = 25$): (a) DFMDock energy, (b) Rosetta energy, and (c) model-derived confidence score. Each point represents a decoy, colored by DockQ-based structural quality: gray (incorrect, DockQ $< 0.23$), orange (acceptable, $0.23 \leq$ DockQ $< 0.49$), blue (medium, $0.49 \leq$ DockQ $< 0.80$), and red (high, DockQ $\geq 0.80$). The vertical dashed line marks the 5 Å threshold for acceptable predictions. The $P_{\text{near}}$ value shown in each panel quantifies the degree of energy funneling (higher is better).

### References

[1] Gaurav Bhardwaj, Vikram Khipple Mulligan, Christopher D Bahl, Jason M Gilmore, Peta J Harvey, Olivier Cheneval, Garry W Buchko, Surya VSRK Pulavarti, Quentin Kaas, Alexander Eletsky, et al. Accurate de novo design of hyperstable constrained peptides. *Nature*, 538(7625): 329–335, 2016.
